# The Genetic Architecture of Gene Expression Levels in Wild Baboons

**DOI:** 10.1101/008490

**Authors:** Jenny Tung, Xiang Zhou, Susan C Alberts, Matthew Stephens, Yoav Gilad

**Affiliations:** Department of Human Genetics, University of Chicago, Chicago, IL, USA; Department of Evolutionary Anthropology and Duke Population Research Institute, Duke University, Durham, NC, USA; Institute of Primate Research, National Museums of Kenya, Nairobi, Kenya; Department of Statistics, University of Chicago, Chicago, IL, USA; Department of Biostatistics, University of Michigan, Ann Arbor, MI, USA; Department of Biology, Duke University, Durham, NC, USA

## Abstract

Gene expression variation is well documented in human populations and its genetic architecture has been extensively explored. However, we still know little about the genetic architecture of gene expression variation in other species, particularly our closest living relatives, the nonhuman primates. To address this gap, we performed an RNA sequencing (RNA-seq)-based study of 63 wild baboons, members of the intensively studied Amboseli baboon population in Kenya. Our study design allowed us to measure gene expression levels and identify genetic variants using the same data set, enabling us to perform complementary mapping of putative *cis*-acting expression quantitative trait loci (eQTL) and measurements of allele-specific expression (ASE) levels. We discovered substantial evidence for genetic effects on gene expression levels in this population. Surprisingly, we found more power to detect individual eQTL in the baboons relative to a HapMap human data set of comparable size, probably as a result of greater genetic variation, enrichment of SNPs with high minor allele frequencies, and longer-range linkage disequilibrium in the baboons. eQTL were most likely to be identified for lineage-specific, rapidly evolving genes. Interestingly, genes with eQTL significantly overlapped between the baboon and human data sets, suggesting that some genes may tolerate more genetic perturbation than others, and that this property may be conserved across species. Finally, we used a Bayesian sparse linear mixed model to partition genetic, demographic, and early environmental contributions to variation in gene expression levels. We found a strong genetic contribution to gene expression levels for almost all genes, while individual demographic and environmental effects tended to be more modest. Together, our results establish the feasibility of eQTL mapping using RNA-seq data alone, and act as an important first step towards understanding the genetic architecture of gene expression variation in nonhuman primates.

## INTRODUCTION

Gene regulatory variation has been shown to make fundamental contributions to phenotypic variation in every species examined to date. This relationship has been demonstrated most clearly at the level of gene expression, which captures the integrated output of a large suite of other regulatory mechanisms. Variation in gene expression levels has been linked to fitness-related morphological, physiological, and behavioral variation in both lab settings and natural populations (e.g., [1–4]; reviewed in [5]), and is a robust biomarker of disease in humans (e.g, [6,7]). In addition, patterns of gene expression are often associated with signatures of natural selection [8–11], suggesting their functional importance even when their phenotypic significance remains unknown.

In primates, the majority of research on the evolution of gene expression has concentrated on cross-species comparisons, particularly using humans, chimpanzees, and rhesus macaques [8,12-22]. Such studies have been important for identifying patterns of constraint on gene expression phenotypes over long evolutionary time scales, and for suggesting candidate loci that might contribute to phenotypic uniqueness in humans or other species. For example, gene expression patterns associated with neurological development appear to have experienced an accelerated rate of change in primates relative to other mammals, with axonogenesis-related and cell adhesion-related genes accelerated specifically in the human lineage [15]. Similarly, differentially expressed genes in human liver are enriched for metabolic function [8], suggesting a potential molecular basis for arguments implicating dietary shifts in the emergence of modern humans [23–25].

Adaptively relevant changes in gene expression levels across species implicate selection on gene expression phenotypes within species, and particularly within populations, the basic unit of evolutionary change. However, in contrast to cross-species comparisons, we still know little about the genetic architecture of gene expression levels in natural nonhuman primate populations. No estimates of the heritability of gene expression traits are available, even for populations that have been intensively studied for many decades. We also do not know whether segregating genetic variation that affects gene expression is common or rare, how the effect sizes of such variants are distributed, or whether they carry a signature indicative of natural selection. If gene regulatory variation has indeed been key to primate evolution, as classic arguments suggest [26], then large gaps therefore remain in our understanding of this process.

Three primary reasons combine to account for the absence of such data. First, until relatively recently, the only feasible approach for measuring genome-wide gene expression levels on a population scale was microarray technology. This constraint limited the diversity of systems that could be assessed because cost-effective, commercially available arrays have only been developed for a handful of taxa. Second, genomic resources, especially detailed catalogues of known genetic variants (e.g., [27,28]), are also limited to a small set of species. The lack of such resources creates major barriers to genome-scale studies of the genetics of gene expression in other organisms, which rely on complementary gene expression and genotype data. Finally, for many taxa, samples suitable for gene expression profiling can be challenging to collect. In nonhuman primates, for example, RNA samples are rarely available even for the most intensively studied natural populations.

Recently, sequencing-based methods for measuring gene expression levels (e.g., RNA-seq) have eliminated the need for species-specific arrays. Comparative genomic studies using RNA-seq have thus vastly expanded the set of taxa for which genome-wide expression data are available (including primates: [15,22]). Importantly, because fragments of expressed genes are resequenced many times in RNA-seq studies, data on genetic variation are also generated in the process. Although these data can be affected by technical biases, several studies have demonstrated the generally high reliability of genotypes inferred from RNA-seq reads [22,29]. Such data can provide important insight into genetic diversity in species for which little other information exists [22]. Additionally, they provide the two ingredients necessary for mapping gene expression traits to genotype, at moderate cost and without the requirement for previously ascertained genetic variants.

Here, we evaluate the potential for such work in an intensively studied wild primate population, the baboons (*Papio cynocephalus*) of the Amboseli basin in Kenya. Forty-three years of prior research on this population have established it as an important model for human social behavior, health, and aging [30], and have facilitated the development of protocols for collecting samples appropriate for gene expression analysis [31–33]. We generated RNA-seq data for 63 individually recognized members of the Amboseli study population. We used these data to explore the frequency, impact, and potential selective relevance of variants associated with variation in gene expression levels, using complementary expression quantitative trait locus (eQTL) mapping and allele-specific expression (ASE) approaches. We found evidence for abundant functional regulatory variation in the Amboseli baboons, and a surprising amount of power to detect these variants even with a modest sample size. We also found that functional variants are depleted in highly conserved genes, consistent with constraint on gene expression patterns. However, among genes with eQTL, we did not find strong support for a relationship between effect size and minor allele frequency. Such a relationship would be consistent with pervasive negative selection on gene expression phenotypes (i.e., selection against variants that produce large perturbations in gene expression levels) and has been suggested by work in humans [34]. Finally, we used our data set to provide the first estimates of the heritability of gene expression levels in wild primates, including the relative contributions of *cis*-acting and *trans*-acting genetic variation.

## RESULTS

### Functional regulatory variation is common in the Amboseli baboons

We obtained blood samples from 63 individually recognized adult baboons in the Amboseli population (Figure 1). From these samples, we produced a total of 1.89 billion RNA-seq reads (mean of 30.0 ± 4.5 s.d. million reads per individual: Table S1). On average, 67.2% of reads mapped to the most recent release of the baboon genome (*Panu2*.*0*), 69.2% of which could be assigned to a unique location. We used the set of uniquely mapped reads to estimate gene-wise gene expression levels for NCBI-annotated baboon RefSeq genes. After subsequent read processing and normalization steps (Methods, Figure S1-S2), we considered variation in gene expression levels for 10,409 genes expressed in whole blood (i.e., all genes for which we could test for *cis*-acting genetic effects on gene expression).

**Figure 1.**
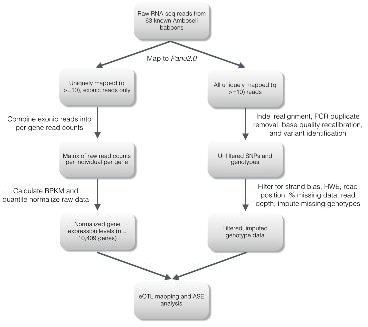
Workflow for identification of functional *cis*-regulatory variants using RNA-seq data.

We also used the RNA-seq reads to identify segregating genetic variants in the Amboseli population. We considered only high confidence sites that were variable within the Amboseli population (Methods and Text S1; Figure S3). As expected [29], these sites were highly enriched in annotated gene bodies (Figure 2; Figure S4). Based on parallel analyses applied to human RNA-seq data, we estimated approximately 97% of these sites to be true positives, and a median correlation between true genotypes and inferred genotypes of 98.7% (Text S2; Figure S5-S6). To identify putative expression quantitative trait loci (eQTL), we focused on variants that passed quality control filters, within 200 kb of the gene of interest. Such variants represent likely *cis*-acting eQTL, which are more readily identifiable in small sample sizes than *trans*-eQTL. To identify cases of allele-specific expression, which provide independent but complementary evidence for functional *cis*-regulatory variation, we focused on genes for which multiple heterozygotes were identified for variants in the exonic regions of expressed genes. We also required a minimum total read depth at exonic heterozygous sites of 300 reads (which should provide high power to detect modest ASE: [35]), resulting in a total set of 2,280 genes tested for ASE.

**Figure 2.**
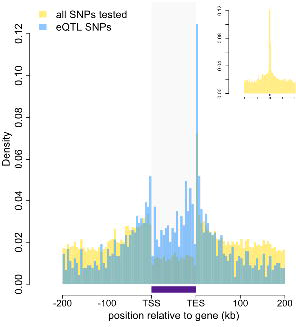
Baboon eQTLs are enriched in and near genes. The locations of all SNPs tested in the eQTL analysis are shown in gold relative to the 5’ most gene transcription start site (TSS) and the 3’ most gene transcription end site (TES) for all 10,409 genes. SNPs detected as eQTL are overplotted in blue, and are enriched, relative to all SNPs tested, near transcription start sites, transcription end sites, and within gene bodies. Gray shaded rectangle denotes the region bounded by the TSS and TES, with gene lengths divided into 20 bins for visibility (because the gene body is thus artificially enlarged, SNP density within genes cannot be directly compared with SNP density outside of genes). Note that SNPs that fall outside of one focal gene may fall within the boundaries of other genes. Inset: Distribution of all SNPs tested relative to the location of genes, highlighting the concentration of SNPs in genes (the peak at the center of the plot).

Both analyses converged to reveal extensive segregating genetic variation affecting gene expression levels in the Amboseli population. At a 10% false discovery rate, we identified eQTL for 1,787 (17.2%) of the genes we analyzed, and evidence for ASE for 510 (23.4%) of tested genes. Consistent with reports in humans (e.g., [36,37]), eQTL were strongly enriched near gene transcription start sites and in gene bodies (Figure 2; controlling for the background distribution of sites tested, which were also enriched in and around genes). Within gene bodies, eQTL were particularly likely to be detected near transcription end sites; this potentially reflects enrichment in 3’ untranslated regions, which are poorly annotated in baboon. Also as expected, genes with eQTL were more likely to exhibit significant ASE and vice-versa (hypergeometric test: p < 10^−25^; Figure 3a). The magnitude and direction of ASE and eQTL were significantly correlated when an eQTL SNP could also be assessed for ASE (n = 123 genes; *r* = 0.719, p < 10^−20^, Figure 3b), and when ASE SNPs were assessed as eQTL (n = 510 genes; *r* = 0.575, p < 10^−45^, Figure 3c).

**Figure 3.**
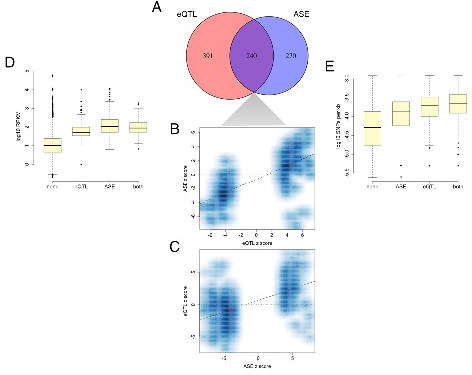
Agreement between eQTL and ASE approaches for identifying functional variants. **(A)** Venn diagram depicting the overlap between genes with significant eQTL and ASE, among genes tested in both cases. Genes with significant eQTL are more likely to have significantly detectable ASE and vice-versa (p = 6.02 x 10^−27^). **(B)** eQTL SNPs in exonic regions that could also be tested for ASE reveal correlated effect sizes (n = 123; p < 7.47 x 10^−21^). **(C)** Similarly, ASE SNPs exhibit effect sizes that are correlated with evidence for eQTL at the same sites (n = 510; p < 2.60 x 10^−46^). **(D)** Detection of ASE is favored for genes with higher expression levels (p = 3.99 x 10^−209^), **(E)** whereas detection of eQTL is favored for genes with greater *cis*-regulatory SNP density (p = 1.05 x 10^−73^).

Combined, the two approaches yielded more information about genetic effects on gene expression than either approach alone. A total of 2,057 genes were associated with either an eQTL or significant ASE, with 240 genes common to both sets (of the 2,280 genes tested in both analyses, 901 of which were associated with an eQTL or significant ASE). Further, the set of genes for which we detected ASE but not a significant eQTL (when tested for both; n = 270) were associated with significantly stronger evidence for harboring an eQTL than genes without ASE (p = 1.7 x 10^−7^), suggesting that these genes likely have moderate effect eQTL that were undetectable given our sample size. The 391 genes identified only in the eQTL analysis were similarly enriched for evidence of ASE (p = 2.4 x 10^−4^). Further investigation revealed that whereas detection of ASE was most strongly favored for highly expressed genes (i.e., higher RPKM: Wilcoxon test: p < 10^−208^; Figure 3d), detection of eQTL was most strongly favored for genes with high local SNP density (p < 10^−72^; Figure 3e). This pattern likely emerges because power to detect ASE is dependent on sequencing read coverage at heterozygous sites [35], which scales with gene expression level. In contrast, power to detect eQTL is affected by the likelihood of performing a test for association against at least one SNP in linkage disequilibrium with a causal variant(s). Functional regulatory variants were thus least likely to be detected for lowly expressed genes with low genetic diversity, potentially reflecting selective constraint on these genes as well as reduced power to identify genetic variants, test for ASE, and map eQTL.

### Increased power to detect eQTL in baboons relative to humans

The number and effect sizes of the eQTL we detected indicate that our power to detect eQTL in the Amboseli population was surprisingly high, especially given that our genotyping data set was limited only to those sites represented in RNA-seq data (i.e., primarily within transcribed regions of moderately to highly expressed genes). Further, while thousands of *cis*-eQTL have been mapped in single human populations, doing so has generally required sample sizes several fold larger than ours [34,38].

To provide a more informative estimate of the difference in power to detect eQTL in baboons relative to humans, we applied the same mapping, data processing, variant calling, and eQTL modeling pipeline to a similarly sized RNA-seq data set on 69 Yoruba (YRI) HapMap samples, in which samples were sequenced to a similar depth [36]. Using our approach for estimating and modeling the gene expression data, but obtaining the genotype data from an independent array platform, we could identify 700 genes with significant eQTL in the YRI data set at a 10% FDR. Approximately half (51%) could be recovered if we only focused on SNPs in transcribed regions. This number (n = 357) therefore reflects the likely theoretical limit of detection for performing eQTL mapping in which SNPs are called based on RNA-seq data (that is, when sequence coverage considerations are not taken into account). Indeed, when eQTL mapping for the YRI was conducted using genotype data obtained from RNA-seq reads (i.e., the same pipeline used for the baboons), we identified 290 genes with eQTL (41.4% of those identified using independently collected genotype data). The RNA-seq-based pipeline therefore reduces the number of genes with detectable eQTL by 50 – 60%. Extrapolation of this estimate suggests that, if genotyping array data had been available for the baboons, we might have identified eQTL for ~3500 - 4000 genes, comparable to results from human data sets with more than 350 samples [38]. To better understand the reasons behind this difference, we investigated three possible explanations.

#### Shifts in the minor allele frequency spectrum

We observed that the minor allele frequency (MAF) spectrum of variants called in the baboon data set included proportionally more intermediate frequency variants and proportionally fewer low frequency variants than in the human data set (Figure 4a, inset). To investigate the degree to which this shift conferred greater power to detect eQTL in the baboons, we simulated eQTL for 10% of the genes in the study by randomly choosing a SNP near each of these genes.

**Figure 4.**
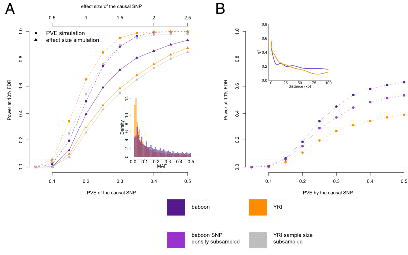
Power to detect eQTL in the Amboseli baboons compared to the HapMap YRI population. **(A)** Simulated eQTL data sets demonstrate that the baboon data set has greater power to detect eQTL (at a 10% FDR threshold) when eQTL are simulated based on effect size (solid lines and triangles) but not when eQTL are simulated based on proportion of variance in gene expression levels explained (PVE: dashed lines and circles). This result likely stems from differences in the minor allele frequency (MAF) spectrum between baboons and YRI (inset), which favors eQTL mapping in the baboons; simulations based on effect size are sensitive to MAF, but simulations based on PVE are not. **(B)** Masking the simulated eQTL SNP demonstrates that the baboon data set has greater power to detect eQTL due to both increased *cis*-regulatory SNP density and more extended LD (inset). Subsampling the SNP density in the baboon data set to the level of the YRI data set reduces the difference in power but does not remove it completely. In B, all results are shown for PVE-based simulations to exclude the effects of the MAF.

We did so in two ways. First, we simulated the effect size of the eQTL, with possible effect sizes ranging from 0.25 to 2.5, in intervals of 0.25 (effect sizes are relative to a standard normal distribution). The power to detect an eQTL of a given effect size is contingent on the relative representation of different genotype classes in a population, and hence MAF (larger MAFs produce a more balanced set of alternative genotypes, and thus more power). Second, we simulated the proportion of variance in gene expression levels (PVE) explained by the eQTL, with possible PVE values ranging from 5% to 50%, in intervals of 5%. In this case, power to detect an eQTL does not depend on MAF because simulating the PVE directly integrates across the combined impact of effect size and MAF (a simulated high PVE eQTL with low MAF implies a large effect size variant). Thus, the impact of the MAF spectrum on the power to detect eQTL is reflected in the differences in power between the baboon data set and the YRI data set in the effect size-based versus the PVE-based simulations. In all cases, we calculated power as the proportion of genes with simulated eQTL recovered at a 10% FDR.

In PVE-based simulations, power to detect simulated eQTL was greater in the YRI data set (Figure 4a, dashed orange line vs dashed purple line), although this advantage disappeared when the YRI data set was subsampled to the same size as the baboon data set (Figure 4a, dashed gray line). However, the baboon data set provided more power to detect eQTL than the YRI data set (whether subsampled or not) when simulations were based on effect size, where power scales with MAF (Figure 4a, solid lines). Based on these differences, we estimate that the power to identify an eQTL of effect size equal to the mean estimated beta in baboons (0.96), is increased in the Amboseli baboons by approximately 1.34-fold (Figure 4a, solid purple line vs solid orange line) as a function of differences in the MAF spectrum alone.

#### Differences in genetic diversity and linkage disequilibrium

Because our RNA-seq-based approach does not identify variants outside of transcribed regions, causal SNPs were probably often not typed. To quantify the power to detect eQTL under this scenario, we again simulated eQTL among genes in the baboon and YRI data sets, but masked the causal sites. Doing so revealed much greater power to identify eQTL in baboons than in humans, across all values of simulated PVE or effect size (Figure 4b; Figure S7). One possible explanation for this observation stems from increased genetic diversity in the baboons compared to the YRI. Indeed, in baboons we tested an average of 45.4 (± 57.0 s.d.) genetic variants for each gene, whereas applying the same pipeline in YRI yielded an average of 20.3 (± 21.4 s.d.) testable variants per gene. An alternative explanation relates to patterns of LD, which we estimate to decay somewhat more slowly in the baboons (Figure 4b, inset). Higher SNP density in baboons increases the likelihood that, when a causal SNP is not typed, a nearby SNP will be available that tags it. Longer range LD suggests that a given SNP could also tag distant causal variants more effectively.

To assess the contributions of SNP density and LD, we refined our simulations by first thinning the SNP density in the baboons to match SNP density in the YRI, and again masking the simulated causal eQTL. As expected, reducing genetic diversity in the baboons reduced the power to detect genes with a true eQTL (Figure 4b, purple dashed line vs pink dashed line). However, it did not completely account for the difference between the human population and the baboon population, suggesting that LD patterns probably contribute to higher eQTL mapping power in baboons as well as SNP density. Specifically, for an eQTL that explains 28% of the variance in gene expression levels (the mean PVE detected in baboons for genes with significant eQTL), we estimate that SNP density and LD effects increase power by 1.21-fold (Figure 4b, purple dashed line vs pink dashed line) and 1.43-fold (Figure 4b, pink dashed line vs orange dashed line), respectively, when causal SNPs are not typed.

Together, our simulations suggest that the MAF spectrum, genetic diversity, and LD patterns increase the number of genes with detectable eQTL in baboons versus the YRI by 2.35-fold overall (1.34x from the MAF, 1.21x from SNP density effects, and 1.43x from LD effects). Further, considering that the effect size estimates in baboons tended to be larger than in the YRI (mean of 0.96 in baboons versus mean of 0.80 in YRI), the actual fold increase estimated from simulations is approximately 6-fold (Figure S7: ratio of purple versus orange lines at these effect sizes). This estimate is remarkably consistent with empirical results from our comparison of the real baboon and YRI data, in which we identified 6.16-fold the number of eQTL in the baboons.

### Mixed evidence for natural selection on gene expression levels

Interestingly, we found that genes harboring eQTL in baboons were also more likely to have detectable eQTL in the YRI (hypergeometric test, p = 2.39 x 10^−7^). Given the sample size limitations of the data sets we considered, this overlap suggests that large effect eQTL tend to be nonrandomly concentrated in specific gene orthologues. This pattern could arise if the regulation of some genes has been selectively constrained over long periods of evolutionary time, whereas others have been more permissible to genetic perturbation. Indeed, we found that the mean per-gene phyloP score calculated based on a 46-way primate comparison was significantly reduced (reflecting less conservation) for genes with detectable eQTL in both species, and greatest for genes in which eQTL were not detected in either case (p < 10^−53^; Figure 5a). We obtained similar results using phyloP scores based on a 100-way vertebrate comparison (p < 10^−21^; Figure S8).

**Figure 5.**
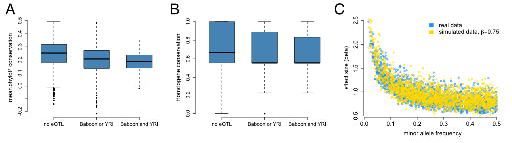
Mixed evidence for negative selection on variants affecting gene expression level. **(A)** Genes that harbor detectable eQTL in baboons, the YRI, or both are more likely to be conserved across long stretches of evolutionary time, based on mean phyloP scores in a 46-way primate genome comparison. **(B)** These genes are also more likely to be lineage specific, based on Homologene annotations. **(C)** Although we detect a strong negative correlation between eQTL effect size and eQTL minor allele frequency, in support of pervasive selection against alleles with large effects on gene expression levels, this correlation also appears when simulating constant eQTL effect sizes, suggesting winner’s curse effects.

eQTL were more likely to be identified for genes with higher genetic diversity (Figure 3), which may account for the relationship between phyloP score and eQTL across species: highly conserved genes are less likely to contain many variable sites. However, genes with eQTL in both species were also less likely to have orthologues in deeply diverged species, based on conservation in Homologene (*β* = −0.036, p = 1.78 x 10^−8^; Figure 5b). Genetic diversity within the baboons is very weakly correlated with Homologene conservation (r^2^ = 0.004). Thus, sequence-level conservation scores and depth of homology across species combine to suggest that eQTL—or at least those with relatively large effect sizes—are least likely to be detected for strongly conserved loci, and most likely to be detected for lineage-specific, rapidly evolving genes. Consistent with this idea, genes involved in basic cellular metabolic processes were under-enriched among the set of genes with eQTL in both species, and enriched among the set of genes for which no eQTL were detected in either species (Tables S2-S3). The set of genes with eQTL in either or both species, on the other hand, were enriched for loci involved in antigen processing, catalytic activity, and interaction with the extracellular environment (e.g., receptors, membrane-associated proteins).

Widespread selective constraint on gene expression levels has been suggested in previous eQTL analyses in humans, with evidence supplied by a strong negative correlation between minor allele frequency and eQTL effect size [34]. This pattern could arise if selection acts against large genetic perturbations, such that variants of large effect would be present only at low frequencies. Consistent with this idea, plotting eQTL effect size versus MAF in the baboons results in a very strong, highly significant negative correlation (r = −0.723, p < 10^−280^; Figure 5c), with no large effect eQTL detected at higher MAFs. However, such a relationship could also be a consequence of the so-called winner’s curse (in which sampling variance leads to upwardly biased effect size estimates: [39]) because the degree of bias in effect size estimation is itself negatively correlated with MAF. Indeed, when we simulated sets of eQTL with constant small effect sizes (*β* = 0.75, close to the mean effect size detected for SNPs with MAF ≥ 0.4), we found that the relationship between estimated effect size and MAF among detected eQTL almost perfectly recapitulated the observed negative correlation. Hence, the correlation between estimated eQTL effect size and MAF in the baboons does not provide strong support for widespread negative selection on gene expression phenotypes within species.

### Genetic and environmental contributions to gene expression variation in wild baboons

Finally, we took advantage of our data set to generate the first estimates of genetic, demographic (age and sex), and environmental contributions to gene expression variation in wild nonhuman primates (Table S4). While our limited sample size leads to high variance around estimates for any individual gene, the median estimates across genes should be unbiased [40], so we concentrated on these overarching patterns. We focused specifically on three social environmental variables of known importance in this population, all of which have been extensively investigated as models for human social environments. These were: i) early life social status, which predicts growth and maturation rates [41,42]; ii) maternal social connectedness to other females, which predicts both adult lifespan and the survival of a female’s infants [43–46]; and iii) maternal social connectedness to males, based on recent evidence that heterosexual relationships have strong effects on survival as well [43].

Overall, we found that genetic effects on gene expression levels tended to be far more pervasive than demographic and environmental effects. Specifically, the median additive genetic PVE was 28.4%, similar to, or slightly greater than, estimates from human populations [47–51]. We applied a Bayesian sparse linear mixed model (BSLMM: [40]) to further partition this additive genetic PVE into two components: a component attributable to *cis*-SNPs (here, all SNPs within 200 kb of a gene) and a component attributable to *trans*-SNPs (all other sites in the genome). Again similar to humans [50,51], we found that more of the additive genetic PVE is explained by the *trans* component (median PVE = 23.8%) than the *cis* component (median PVE = 2.9%) (Figure 6). Unsurprisingly, we estimated a larger *cis*-acting component for genes in which functional *cis*-regulatory variation was detected in our previous analysis (median PVE = 10.2% among eQTL genes and median PVE = 5.0% among ASE genes).

**Figure 6.**
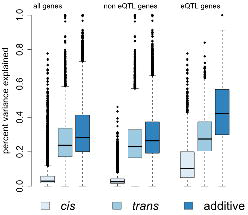
Genetic contributions to variance in gene expression levels in wild baboons. Proportion of variance in gene expression levels estimated for all genes, genes without detectable eQTL, and genes with detectable eQTL. Additive genetic effects on gene expression variation, especially *cis*-acting effects, are larger for eQTL genes than for other genes.

In contrast to the substantial genetic effects we detected, the median PVE explained by age and sex were 1.89% and 0.82% respectively (Figure S9). The distribution of PVE explained by age was significantly greater than expected by chance (Kolmogorov-Smirnov test on binned PVEs, in comparison to permuted data: p < 10^−11^), whereas that explained by sex was not (p = 0.100); large sex effects tended to be constrained to a small set of genes on the X chromosome (Figure S9). Of the early environmental variables we investigated, only maternal social connectedness to males explained more variance in gene expression levels than expected by chance (p = 4.19 x 10^−3^), with a median PVE of 1.9%. Notably, while social connectedness to males (i.e., heterosexual bonds) and social connectedness to females (i.e., same-sex bonds) are both known predictors of longevity in the Amboseli baboons, previous analyses suggest that their effects are largely independent [43]. Our result extends this observation to the early life effects of maternal social connectedness on variance in gene expression levels.

Taken together, our data suggest that while almost all genes are influenced by genetic variation, the effects of demographic and environmental parameters generally modest for any single aspect of the environment. However, in at least some cases, we find evidence that early environmental effects on gene expression levels appear to persist across the life course, as has previously been demonstrated in laboratory settings and in response to severe early adversity in humans (e.g., [52–54]).

## DISCUSSION

Much of what we know about genetic contributions to variation in gene expression levels in primates (and vertebrates more generally) comes from the extensive body of research on humans. However, increasing evidence indicates that humans are demographically unusual: compared to other primates, humans exhibit low levels of neutral genetic diversity and a low long-term effective population size [22,55,56]. Further, humans are distinguished from other primates by recent explosive population growth [57,58]. While late Pleistocene population expansion has been suggested for some nonhuman primates, including chimpanzees and Chinese-origin rhesus macaques [56,59], none have undergone the extreme levels of population increase that characterized humans. Indeed, evidence from microsatellite data suggests that the long-term effective population size of baboons actually may have contracted during this period [60].

These differences are not simply of historical interest, but also important for understanding the genetic architecture of traits measured in the present day. Differences in demographic history not only affect overall levels of genetic variation and the minor allele frequency spectrum, but also the mean effect size of sites that contribute to phenotypic variation [61]. Interestingly, demographic history does not impact overall trait heritability [61,62], perhaps explaining why we estimated mean additive genetic PVEs for gene expression levels in baboons that are similar to those estimated for humans. However, demographic history can influence the power to detect individual genetic contributions to phenotypic variation. Large-scale population expansion of the type that occurred in human history appears to reduce power to identify genotype-phenotype correlations for fitness-related traits [61]. This observation may account, in part, for our ability to identify many more functional regulatory variants in the baboons than we expected based on previous studies in humans.

However, while our analysis extends previous observations that large effect eQTL are non-randomly distributed, we found mixed evidence for widespread negative selection on gene expression levels. Specifically, within the baboons alone, we found that the negative relationship between eQTL effect size and minor allele frequency was explicable based on winner’s curse effects alone. Thus, increased power to identify functional regulatory variants in the baboons is probably not due to pervasive associations between gene expression levels and fitness. In contrast, stronger evidence for selection on gene expression patterns stems from our cross-species comparisons. In particular, we observed that genes with eQTL in baboons significantly overlapped with genes with eQTL in humans, and that these genes as a class also tended to be less constrained at the sequence level. This result suggests that genes vary in their tolerance of functional regulatory genetic variation, and, intriguingly, that gene-specific robustness to genetic perturbation may be a conserved property across species.

Because no comparable data are yet available for other large mammal populations, including for other baboons, it is unclear whether our results are typical or instead a consequence of the Amboseli population’s own unique history. In particular, the population has experienced recent admixture between yellow baboons, the dominant taxon, and closely related anubis baboons (*Papio anubis*) [63,64]. Admixture, which appears to be relatively common in natural populations [65], can have important consequences for genetic diversity and LD patterns, and may have contributed to our ability to readily map gene expression phenotypes. Comparison to a non-admixed baboon population could help resolve this question. More generally, our results encourage further investigation of the relationship between demography and trait genetic architecture in other populations, as has been suggested for humans [61] but could also be profitably extended to nonhuman model systems. Such comparisons would provide an empirical basis for testing predicted relationships between demographic history and the power to identify genotype-phenotype associations. From an applied perspective, they could also help identify animal models that favor more highly powered association mapping studies, a strategy that has already been heavily exploited in domestic dogs [66,67] and suggested for rhesus macaques [56]. While the same sites will probably rarely be associated with the same traits across species, this strategy could help identify molecular mechanisms that are conserved across humans and animal models (e.g., [68]).

Finally, our data—the first profile of genome-wide gene expression levels in a wild primate population—serve as a useful proof of principle of the ability to concurrently generate genome-wide gene expression phenotype and genotype data, and to relate them to each other using eQTL and ASE approaches. Intensively studied natural primate populations—some of which have been studied continuously for thirty or more years—have emerged as important phenotypic models for human behavior, health, and aging. The approach we used here provides a way to leverage these models for complementary genetic studies as well, especially if eQTL prove to be strongly enriched for sites associated with other traits, as in humans [69]. Although preliminary, our results highlight the increasing feasibility of integrating functional genomic data with phenotypic data on known individuals in the wild. For example, our data set revealed a number of genes in which variation in gene expression levels could be mapped to an identifiable eQTL, validated using an ASE approach, and also linked to early life environmental variation. Such cases suggest the potential for future investigations of the molecular basis of persistent environmental effects, including whether genetic and environmental effects act additively or interact.

## METHODS

### Study subjects and blood sample collection

Study subjects were 63 individually recognized adult members (26 females and 37 males) of the Amboseli baboon population. All study subjects were recognized on sight by observers based on unique physical characteristics. To obtain blood samples for RNA-seq analysis, each baboon was anesthetized with a Telazol-loaded dart using a handheld blowpipe. Study subjects were darted opportunistically between 2009 and 2011, avoiding females with dependent infants and pregnant females beyond the first trimester of pregnancy (female reproductive status is closely monitored in this population, and conception dates can be estimated with a high degree of accuracy). Following anesthetization, animals were quickly transferred to a processing site distant from the rest of the group. Blood samples for RNA-seq analysis were collected by drawing 2.5 mL of whole blood into PaxGene Vacutainer tubes (Qiagen), which contain a lysis buffer that stabilizes RNA for downstream use. Following sample collection, study subjects were allowed to regain consciousness in a covered holding cage until fully recovered from the effects of the anesthetic. They were then released within view of their social group; all subjects promptly rejoined their respective groups upon release, without incident.

Blood samples were stored at approximately 20 C overnight at the field site. Samples were then shipped to Nairobi the next day for storage at −20 C until transport to the United States and subsequent RNA extraction.

### Ethics statement

Samples and data used in this study were collected from wild baboons living in the Amboseli ecosystem of southern Kenya. All behavioral, environmental, and demographic data are gathered as part of noninvasive observational monitoring of known individuals within the study population. This research is conducted under the authority of the Kenya Wildlife Service (KWS), the Kenyan governmental body that oversees wildlife (permit number NCST/5/002/R/777 to SCA and NCST/RCD/12B/012/57 to JT). As the animals are members of a wild population, KWS requires that we do not interfere with injuries to study subjects inflicted by predators, conspecifics, or through other naturally occurring events (e.g., falling out of trees). To collect blood samples, we perform temporary immobilizations using the anesthetic Telazol, delivered via a handheld blowgun. Permission to perform this procedure was granted through KWS, and was performed under the supervision of a KWS-approved Kenyan veterinarian, who monitored anesthetized animals for hypothermia, hyperthermia, and trauma (no such events occurred during our sample collection efforts). Observational and sample collection protocols were approved though IACUC committees at Duke University (current protocol is A028-12-02 to SCA and JT) and the University of Chicago (ACUP 72080 to YG).

### Gene expression profiling using RNA-seq

For each RNA sample (one per individual), we constructed an RNA-seq library suitable for measuring whole genome gene expression using Dynal bead poly-A mRNA purification and a standard Illumina RNA-seq prep protocol. Each library was randomly assigned to one lane of an Illumina Genome Analyzer II instrument and sequenced to a mean depth of 30 million 76-base pair reads (± 4.5 million reads s.d., Table S1). The resulting reads were mapped to the baboon genome (*Panu2.0*) using the efficient short-read aligner *bwa* [70]. To recover reads that spanned putative exon-exon junctions, and therefore could not be mapped directly to the genome, we used the program *jfinder* on reads that did not initially map [36]. Finally, we filtered the resulting mapped reads data for low quality reads (quality score < 10) and for reads that did not map to a unique position in the genome. To assign reads to genes, we used the RefSeq exon annotations for *Panu* 2.0 (downloaded Sept 6, 2012). In downstream analyses, we considered only highly expressed genes that had non-zero counts in more than 10% individuals, and that had mean read counts greater than or equal to 10 (excluding the gene for beta-globin).

We then performed quantile normalization across samples followed by quantile normalization for each gene individually, resulting in estimates of gene expression levels for each gene that were distributed following a standard normal distribution. This procedure effectively removed GC bias in gene expression level estimates (Figure S2). For eQTL mapping, ASE analysis, and PVE estimation for sex and age we used all 63 individuals. For PVE estimation for maternal rank and social connectedness, missing data meant that we conducted our analysis on n = 52 and n = 47 individuals, respectively.

### Variant identification and genotype calls

To identify genetic variants in the baboon data set, we used the Genome Analysis Toolkit (v. 1.2.6; [71,72]). Because no validated reference set of known genetic variants are available for baboon, we performed an iterative bootstrapping procedure for base quality score recalibration (SI). In the final call set, we removed all sites i) that were monomorphic in the Amboseli samples; ii) for which genotype data were missing for more than 12 individuals (19%) in the data set; iii) that deviated from Hardy-Weinberg equilibrium; and iv) that failed filters for variant confidence, mapping quality, strand bias, and read position bias (Text S1). We then retained sites with a minimum quality score of 100 located within 200 kb of a gene of interest, and that were sequenced at a mean coverage ≥5x across all samples in the data set. We validated our quality control and filtering steps by performing the same procedure on an RNA-seq data set from the HapMap Yoruba population (Text S2). These steps resulted in a set of 64,432 single nucleotide polymorphisms carried forward into downstream analysis (30,938 for the YRI). For eQTL mapping analysis, missing genotypes in this final set were imputed using BEAGLE [73].

To estimate genome-wide LD, we followed the approach of Eberle and colleagues [74], which uses allele frequency-matched SNPs to calculate pair-wise LD. Specifically, we selected SNPs with MAFs greater than 10% and divided them into four subgroups (MAF between 10%-20%; MAF between 20%-30%; MAF between 30%-40%; and MAF between 40%-50%). We then calculated pair-wise *r^2^* for all SNP pairs within 100kb in each subgroup using VCFtools [75] and combined values from all four subgroups.

### eQTL mapping

To identify *cis*-acting eQTLs in the baboon data set, we used the linear mixed model approach implemented in the program GEMMA [76]. This model provides a computationally efficient method for eQTL mapping while explicitly accounting for genetic non-independence within the sample; in our case, some individuals in the data set are related (although overall relatedness was low: the median kinship coefficient across all pairs was 0.015 (mean = 0.024 ± 0.033 s.d.).

For each gene, we considered all variants within 200kb of the gene as candidate eQTLs. For each variant, we fitted the following linear mixed model:

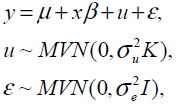

and tested the null hypothesis H_0_: *β* = 0 versus the alternative H_1_: *β* ≠ 0. Here, y is the *n* by 1 vector of gene expression levels for the *n* individuals in the sample; *μ* is the intercept; x is the *n* by 1 vector of genotypes for the variant of interest; and *β* is the variant’s effect size. The *n* by 1 vector of u is a random effects term to control for individual relatedness and other sources of population structure, where the *n* by *n* matrix K=XX^T^/p provides estimates of pairwise relatedness derived from the complete 63 x 64,432 genotype data set X. Residual errors are represented by *ε*, an n by 1 vector, and MVN denotes the multivariate normal distribution.

We took the variant with the best evidence (i.e., lowest p-value) for association with gene expression levels for each gene, and then calculated corrected gene-wise q-values (with a 10% false discovery rate threshold) via comparison to the same values obtained from permuted data (similar to [36,77]). We discuss possibly confounding effects of using RNA-seq data for this analysis in the Supplementary Information (Text S3).

### ASE detection

To identify ASE, we focused on SNPs within gene exons with Phred-scaled quality scores greater than 10. We further required that these sites have more than five reads in more than two individuals and more than 300 total reads across all heterozygous individuals. After these filtering steps, we retained 8,154 SNPs associated with 2,280 genes for ASE analysis.

For each variant, we considered a beta-binomial distribution (following [36]) to model the number of reads from the (+) haplotype (denoted as x_i_^+^) or the number of reads from the (-) haplotype (denoted as x_i_^−^), conditional on the number of total reads (denoted as y_i_=x_i_^+^ + x_i_^−^), for each individual i, or

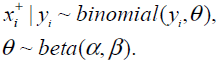

We tested the null hypothesis H_0_: *α* = *β* versus the alternative H_1_: *α* ≠ *β* using a likelihood ratio test, where we maximized the likelihoods in the null and alternative models using the R function *optim*. Again, we took the variant with the lowest p-value for each gene, and then calculated corrected gene-wise q-values (using a 10% false discovery rate threshold) via comparison to the same values obtained from an empirical null distribution. To construct the empirical null distribution, we performed the same analysis after substituting the x_i_^+^ value for each variant of interest, for each heterozygous individual, with a randomly selected x_i_^+^ value from a heterozygous site elsewhere in the genome (contingent on that site having the same number of total reads, y_i_).

### Power simulations

To assess the relative power of eQTL mapping in baboons versus the YRI data set, we randomly selected 10% of the genes in each data set to harbor eQTL. For each of these simulated eQTL genes, we then randomly chose a SNP among all the *cis*-SNPs tested (i.e., all variable sites that passed quality control filters and fell within 200 kb of a gene of interest) and assigned it as a causal eQTL. The impact of the eQTL was simulated using either effect size, in which we simulated a constant effect size between 0.25 and 2.5 (in intervals of 0.25) or PVE, in which we chose an effect size that explained a specific proportion of variance in gene expression levels (from 5% to 50%, in intervals of 5%). We then simulated gene expression levels by adding the effect of the simulated *cis*-eQTL SNP to residual errors drawn from a standard normal distribution. To calculate the FDR, we also simulated a set of genes with no eQTL. For each combination of effect sizes and population (baboon or YRI), and for each simulation scenario (e.g., with the causal SNP masked or unmasked, with SNP density thinned in the baboons, or using PVE versus a constant effect size), we performed 10 replicates. For each replicate, we calculated the power to detect eQTL as the proportion of simulated eQTL genes recovered at a 10% empirical FDR.

### Evidence for patterns consistent with natural selection on gene expression levels

We investigated the relationship between conservation level and the presence of detectable eQTL in the Amboseli baboons or the YRI using phyloP conservation scores [78] and Homologene conservation of orthology across species. For the former, we extracted the per-site phyloP score from the 46-way primate comparison or 100-way vertebrate comparison on the UCSC Genome Browser for each base contained within the annotated exons (including untranslated regions) used for mapping RNA-seq reads in the YRI. We then calculated the average phyloP score across all exons associated with a given gene. We obtained Homologene scores from the CANDID database [79]. In both cases, we used linear models to test for a relationship between conservation level and three categories of genes: those with no detectable eQTL in either the baboons or YRI; those with a detectable eQTL in one of the two species; and those with a detectable eQTL in both species.

To investigate whether the correlation between minor allele frequency and eQTL effect size could be a result of winner’s curse effects, we extracted the results from our simulations in which the causal variant was masked and the true effect size was fixed at a small value (beta = 0.75). We then calculated the correlation between the estimated effect size (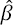) from these simulations against minor allele frequency, for detected eQTL only.

### Estimation of genetic contributions to gene expression

We used the Bayesian sparse linear mixed model (BSLMM) approach implemented in the GEMMA software package [76] to estimate the genetic contribution to gene expression variation. Specifically, for each gene, we fit the following model

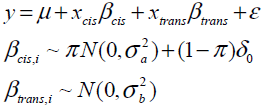

where y is the *n* by 1 vector of gene expression levels for *n* individuals; μ is the intercept; x_cis_ is an n by p_cis_ matrix of genotypes for p_cis_ *cis*-SNPs and β_cis_ are the corresponding effect sizes; x_trans_ is an n by p_trans_ matrix of genotypes for p_trans_ *trans*-SNPs and β_trans_ are the corresponding effect sizes; and *ε* is an n by 1 vector of i.i.d. residual errors. We used different priors for *cis*-acting effects and *trans*-acting effects to capture different properties for the two components. Specifically, the spike-slab prior on the *cis* effects β_cis_ captures our prior belief that only a small proportion of local SNPs have *cis* effects and these effects are relatively large. The normal prior on the *trans* effects captures our prior knowledge that *trans*-acting SNPs tend to be relatively difficult to find and have relatively small effects. In addition, because p_cis_ is small and p_trans_ approximately equals p, the number of total SNPs, we used p instead of p_trans_ to facilitate computation. We used Markov chain Monte Carlo (MCMC) to fit the model with 1000 burn-in and 10000 sampling steps. We obtained posterior samples of β_cis_ and β_trans_ to calculate the PVE attributed by each of the two components, as well as the total additive genetic PVE contributed by both components.

To calculate PVE values for demographic and environmental predictors (described in detail in Text S3), we again used the linear mixed model approach implemented in GEMMA to control for additive genetic effects. For each gene, we fit the following model:

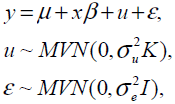

where x is the *n* by 1 vector of values for the demographic or environmental predictor of interest and *β* is its coefficient. The *n* by 1 vector of u is a random effects term with K=XX^T^/p controlling for additive genetic effects. We calculated the PVE estimate as var(x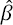)/var(y), where var denotes the sample variance.

## ACKNOWLEDGMENTS

We thank the Kenya Wildlife Services, Institute of Primate Research, National Museums of Kenya, National Council for Science and Technology, members of the Amboseli-Longido pastoralist communities, Tortilis Camp, and Ker & Downey Safaris for their assistance in Kenya. We also thank J. Altmann for general support and access to the Amboseli data set and samples; R.S. Mututua, S. Sayialel, J.K. Warutere, M. Akinyi, T. Wango, and V. Oudu for invaluable assistance with sample collection; K. Michelini for assistance with RNA-seq data generation; S. Mukherjee for advice on statistical analysis; and L.B. Barreiro for comments on an earlier draft of the manuscript. Finally, we thank the Baylor College of Medicine Human Genome Sequencing Center for access to the current version of the baboon genome assembly (*Panu 2.0*).

## SUPPORTING INFORMATION CAPTIONS

**Text S1:** Read mapping and SNP quality filtering pipeline

**Text S2:** Estimated accuracy of SNP genotypes using human RNA-seq data

**Text S3:** Possible confounds associated with eQTL mapping using RNA-seq data

**Text S4:** Demographic and environmental variables

**Table S1:** Read mapping summary

**Table S2:** Gene Ontology analysis for genes with no eQTL in baboon or YRI

**Table S3:** Gene Ontology analysis for genes with eQTL in either or both baboon and YRI

**Table S4:** Demographic and environmental data

**Figure S1:** Detailed workflow for gene expression level estimation

**Figure S2:** Elimination of GC bias via quantile normalization

**Figure S3:** Detailed workflow for SNP genotyping

**Figure S4:** Location of analyzed SNPs relative to genes

**Figure S5:** Accuracy of genotype calls for SNPs independently typed in HapMap3

**Figure S6:** PCA projection of YRI samples using the RNA-seq-based pipeline versus independently typed SNPs

**Figure S7:** Power simulations for masked eQTL based on effect size

**Figure S8:** Correlation between eQTL detection and mean phyloP scores based on 100-way vertebrate comparison

**Figure S9:** PVE explained by demographic and early environmental variables

**Figure S10:** Coverage by genotype call

## ACCESSION NUMBERS

All gene expression data will be deposited into the Gene Expression Omnibus upon publication; raw RNA-seq reads will also be deposited into NCBI’s Short Read Archive.

